# Co-stimulation with opposing macrophage polarization cues leads to orthogonal secretion programs in individual cells

**DOI:** 10.1101/2020.04.04.025536

**Authors:** Andrés R. Muñoz-Rojas, Ilana Kelsey, Jenna Pappalardo, Kathryn Miller-Jensen

## Abstract

Macrophages are innate immune cells that contribute to fighting infections, tissue repair, and maintaining tissue homeostasis. To enable such functional diversity, macrophages resolve potentially conflicting cues in the microenvironment via mechanisms that remain unclear. Here, we used single-cell RNA sequencing to explore how individual macrophages respond when co-stimulated with the inflammatory stimuli, LPS+IFN-γ, and the resolving cytokine, IL-4. We found that co-stimulated macrophages displayed a distinct global transcriptional program. However, variable negative cross-regulation between some LPS+IFN-γ- and IL-4-specific genes resulted in significant cell-to-cell heterogeneity in transcription. Interestingly, negative cross-regulation led to mutually exclusive expression of the T-cell-polarizing cytokines *Il6* and *Il12b* versus the IL-4-associated factors *Arg1* and *Chil3* in single co-stimulated macrophages, and single-cell secretion measurements showed that these specialized functions were maintained for at least 48 hours. Overall, our study suggests that increasing functional diversity in the population is one strategy macrophages use to respond to conflicting environmental cues.

## Introduction

Macrophages respond to a large range of stimuli to aid in development, tissue repair and immunity^1,2^. These polarization responses must be strong enough to defend against pathogens and tissue stress, but sufficiently plastic to accommodate changes in the microenvironment. The M1 inflammatory program, induced as a response to infectious stimuli (e.g. lipopolysaccharide [LPS] and/or interferon-gamma [IFN-γ]) and the M2 program, induced by resolving stimuli (e.g. interleukin-4 [IL-4]), represent two extremes on a spectrum of macrophage responses^3–5^. The M1 polarization program is typically associated with a proinflammatory and anti-bacterial phenotype, while the M2 program is associated with wound-healing, tissue repair and helminth response^5,6^. While this M1-M2 paradigm has been useful in uncovering key regulatory elements in the innate immune response, it is clear that macrophages *in vivo* display a more complex polarization response^7,8^.

There are several possible sources for the complex polarization states observed *in vivo.* First, the vast number of stimuli that can activate macrophages enables these cells to display a large range of functional responses^7,9^. Second, because macrophage polarization does not induce terminal differentiation programs, functional responses by macrophages are plastic and may switch states in response to changing environmental conditions^10^. Additionally, macrophages *in vivo* are constantly exposed to multiple and sometimes conflicting cues. Examples of this include coexisting inflammatory and immunosuppressive signals during the resolution phase of inflammation^11^ or the complex pro- and anti-tumor microenvironments established inside a tumor^12,13^. Together, the flexible and diverse range of macrophage responses and the stimuli they receive result in a complex polarization spectrum observed *in vivo*.

There is extensive crosstalk in the polarization programs in populations of macrophages presented with opposing cues^14,15^; however, the complexity of macrophage polarization has not yet been explored at single-cell resolution. Such single-cell measurements are important because macrophage populations display significant cell-to-cell heterogeneity in their responses even following acute stimulation with LPS^16–18^. The heterogeneity observed *in vitro* is also reflected *in vivo* in numerous studies of macrophage heterogeneity in a variety of disease states, including cancer, liver cirrhosis, and wound healing^19–23^. It is therefore important to characterize how individual macrophages respond to and resolve conflicting cues to coordinate a cohesive immune response within complex tissue microenvironments.

To study individual macrophage responses to opposing polarization cues presented simultaneously, we profiled macrophages stimulated with LPS+IFN-γ, IL-4, or both by single-cell RNA sequencing (scRNA-seq) and by single-cell secretion profiling^17,24^. On a global scale, murine bone marrow-derived macrophages (BMDMs) stimulated with LPS+IFN-γ and IL-4 acquired a unique transcriptional state distinct from the states induced by single stimuli. However, we observed extensive crosstalk between polarization programs in individual macrophages presented with opposing cues, but the extent of this crosstalk varied across cells and with the markers used to define polarization state. Using a combination of neural-networks and statistical analysis, we find a subset of genes from each single-stimulus gene program that are not expressed together in co-stimulated cells, including the T-cell polarizing cytokines *Il6* and *Il12b,* induced by LPS+IFN-γ, and the canonical M2-associated targets *Arg1* and *Chil3,* induced by IL-4. Measurement of highdimensional single-cell secretion profiles confirms that, in most cases, single-cells express either an LPS+IFN-γ-like or IL-4-like secretion phenotype after long-term co-stimulation with opposing cues. Together, our results provide insight into the heterogeneous response of macrophages exposed to opposing polarization cues.

## Results

### Macrophages express a global mixed gene expression program following co-stimulation with LPS+IFN-γ and IL-4

To test how co-stimulation with opposing polarization cues affects the gene expression programs of individual macrophages, we stimulated BMDMs with LPS+IFN-γ, IL-4, or a combination of LPS+IFN-γ and IL-4, and profiled the cells using scRNA-seq (**Fig. 1a**). In cell populations, we observed that in response to 10 ng/ml LPS+10 ng/ml IFN-γ, expression of canonical M1-associated genes (including *Nos2, Tnf, Il6,* and *Il12b*) peaked between 2 and 6 hours, while in response to 100 ng/ml IL-4, expression of canonical M2-associated genes (including *Arg1* and *Chil3)* continued to rise by 8 hours (**Supplementary Fig. 1a**). Therefore, we chose 6 hours as an optimal stimulation time to capture both transcription programs.

**Fig. 1.**
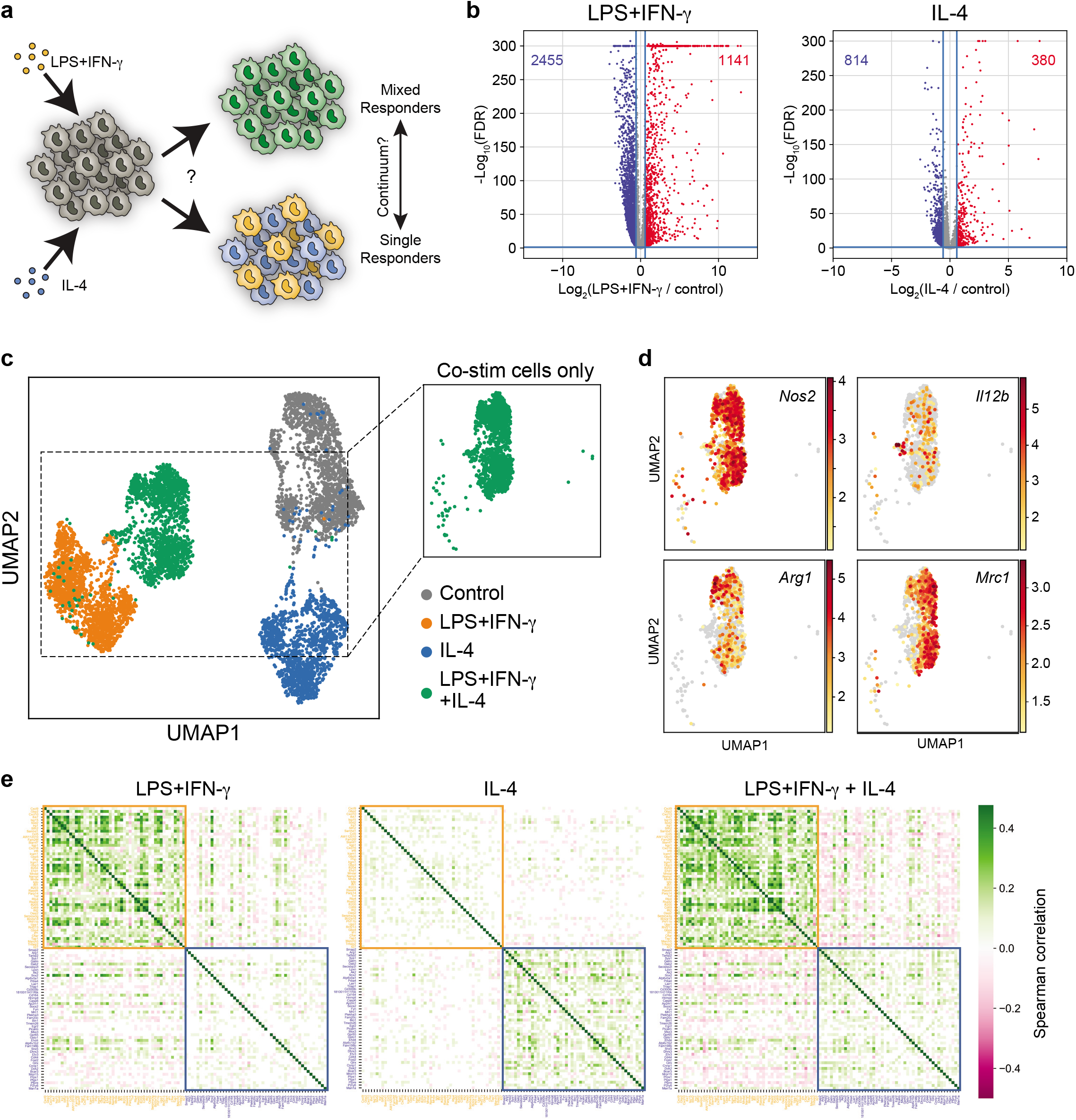
Co-stimulation with LPS+IFN-γ and IL-4 induces a global mixed gene expression program across individual macrophages. **a** Schematic depicting how co-stimulation could lead to either mixed responses or specialized responses in individual macrophages. **b** Volcano plot of differential gene expression after stimulation for 6 h with 10 ng/ml LPS+10 ng/ml IFN-γ (left) or 100 ng/ml IL-4 (right) relative to control. Genes significantly upregulated (red) and downregulated (blue) are identified by a change in expression ≥ 1.5-fold and a false discovery rate (FDR) < 0.05 relative to expression in the untreated condition. FDR calculated by a Wilcoxon rank-sum test with Benjami-ni-Hochberg correction. **c-d** UMAP visualization of single cells from all samples colored by treatment (**c**) or expression intensity for canonical markers of interest (**d**). Color bar indicates gene expression levels, shown as ln(transcript count + 1). **e** Heatmaps of Spearman correlations for the top 50 unique upregulated genes following stimulation with LPS+IFN-γ, IL-4, or both, within single cells from each treatment condition. The top 50 genes upregulated by LPS+IFN-γ or IL-4 are highlighted and boxed in yellow and blue, respectively. Color bar indicates the sign and magnitude of the correlation coefficient. The coefficients for gene pairs with a correlation *p*-value > 0.05 are set to 0.

We also explored a range of LPS and IL-4 dose combinations and observed that co-stimulation with 10 ng/ml LPS+10 ng/ml IFN-γ and 100 ng/ml IL-4 resulted in significant cross-inhibition of LPS+IFN-γ-stimulated IL-6 and IL-12p40 and of IL-4-stimulated *Arg1* and Chil3l3 at 24 hours (**Supplementary Fig. 1b**). In contrast, LPS+IFN-γ-stimulated TNFα and *Nos2* were not significantly inhibited at any dose combination. Thus, we concluded that this dose combination might reveal interesting behaviors in individual cells.

For sc-RNAseq, we profiled ~1,500 single cells per condition to ensure high-quality data with low doublet rates and high sequencing depth per cell (see **Methods**). Our final dataset had an average of 30,262 unique reads per cell and 4,076 genes detected per cell. Stimulation with LPS+IFN-γ upregulated more genes than IL-4 stimulation when compared to unstimulated cells (1141 and 380 genes, respectively), with only 95 of these genes being induced by both conditions (**Fig. 1b**). Interestingly, both LPS+IFN-γ and IL-4 stimulation caused the apparent transcriptional downregulation of thousands of genes normally expressed in unstimulated cells, with the number of downregulated genes substantially outnumbering the number of upregulated genes (**Fig. 1b**).

We performed dimensionality reduction using uniform manifold approximation and projection (UMAP)^25,26^ on the full transcriptional signature to visualize how cells mapped across treatments. We observed that almost all co-stimulated macrophages clustered separately from those stimulated with only one cue and from the unstimulated control population, suggesting that co-stimulation induced a distinct global transcriptional state (**Fig. 1c**). This result also demonstrated that most macrophages were able to respond to both stimuli. This is consistent with the observation that macrophages co-stimulated with LPS+IFN-γ and IL-4 displayed robust Stat1 and Stat6 phosphorylation downstream of the IFN-γ-receptor or IL-4-receptor, respectively, indicating that a majority of macrophages respond to both signals (**Supplementary Fig. 1c**).

Although the transcriptional state of co-stimulated macrophages was distinct from cells treated with LPS+IFN-γ or IL-4 alone, we observed cell separation within the co-stimulation cluster. For example, we observed heterogeneous expression of canonical genes associated with LPS+IFN-γ stimulation, *Nos2* and *Il12b,* versus canonical genes associated with IL-4 stimulation, *Arg1* and *Mrc1* (**Fig. 1d**), suggesting that there is cell-to-cell variability in the extent of each individual cell’s response to LPS+IFN-γ versus IL-4 stimulation. To further explore this possibility, we measured the Spearman correlation between genes uniquely upregulated by either LPS+IFN-γ or IL-4 alone across all single cells to identify which genes are co-expressed or mutually inhibited across single cells. As expected, genes upregulated by LPS+IFN-γ alone were more likely to exhibit positive correlations with other LPS+IFN-γ-induced genes, and either no correlation or negative correlation with IL-4-induced genes (**Fig. 1e**). Similarly, genes upregulated by IL-4 alone were more likely to be positively correlated with other IL-4-induced genes, and uncorrelated or negatively correlated with LPS+IFN-γ-induced genes. Co-stimulated cells generally exhibited weak positive correlations within the LPS+IFN-γ-induced genes and the IL-4-induced genes, while also exhibiting weak negative correlations between some genes across the two programs (**Fig. 1e** and **Supplementary Fig. 1d**). This suggests that on average macrophages expressing core genes of one program exhibited slightly reduced expression of core genes of the other program. Of note, not all correlations between LPS+IFN-γ-induced genes and IL-4-induced genes were negative, indicating that some LPS+IFN-γ-induced genes are co-expressed with IL-4-induced genes. However, the strongest negative correlations were between genes induced by the two opposing stimuli, suggesting that single cells may skew gene expression towards either the LPS+IFN-γ or the IL-4 transcriptional program in response to co-stimulation.

### Co-stimulation with LPS+IFN-γ and IL-4 induces transcriptional cross-regulation that varies substantially across individual cells

We next focused on how co-stimulation with LPS+IFN-γ and IL-4 affected the expression of genes uniquely upregulated by either LPS+IFN-γ or IL-4 alone (**Fig. 2a**), which we refer to as the core gene programs. For both LPS+IFN-γ- and IL-4-induced genes, co-stimulation with the other cue caused both transcriptional upregulation and inhibition in a subset of core genes belonging to each program, consistent with our own and previously reported observations^14^. Among the LPS+IFN-γ-induced core genes, 87 were inhibited by co-stimulation and 97 were augmented by co-stimulation, while among the IL-4-induced core genes, 196 were inhibited and 16 were augmented by co-stimulation (**Fig. 2b**).

**Fig. 2.**
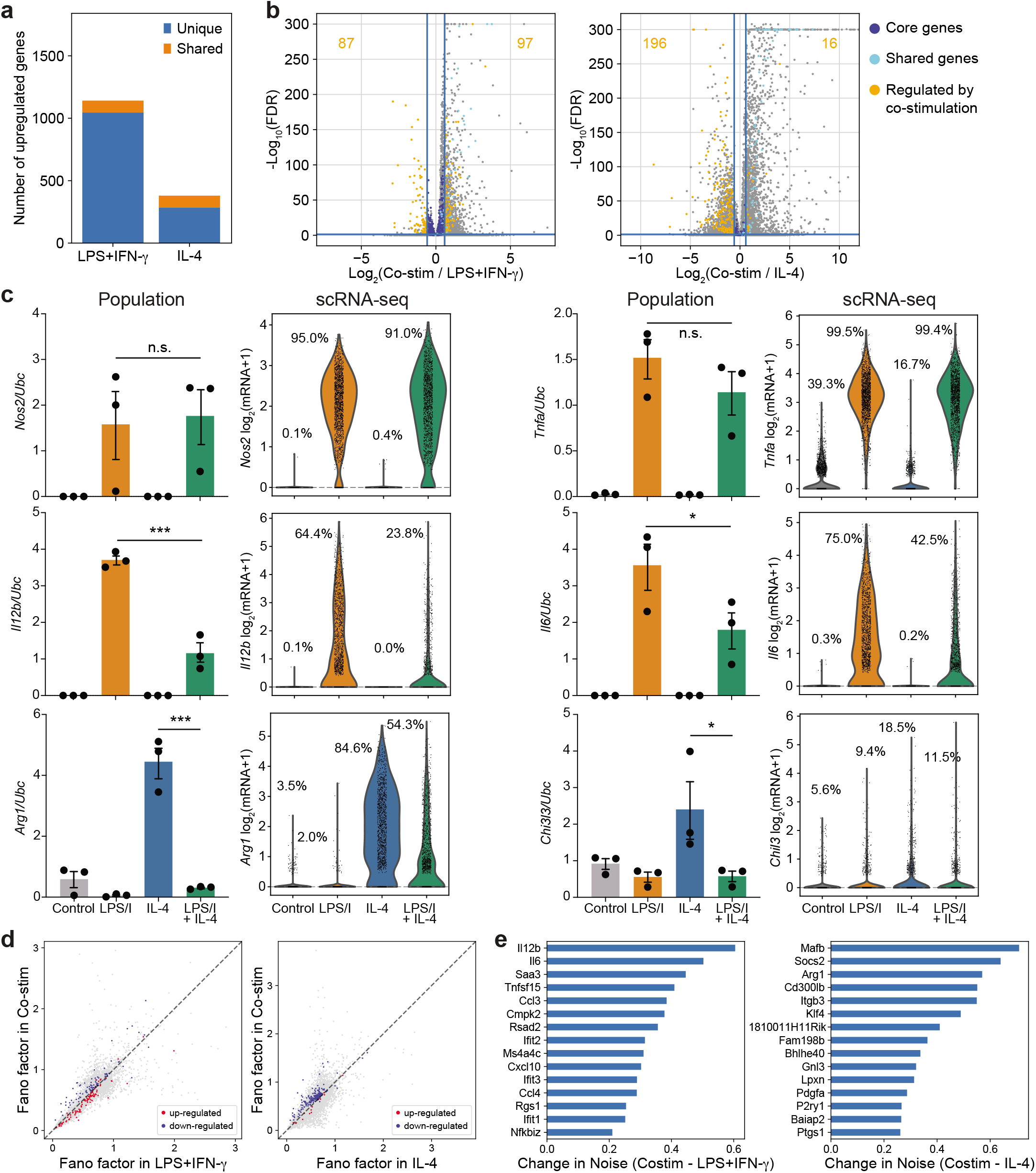
Cross-inhibition of gene expression following LPS+IFN-γ and IL-4 co-stimulation does not occur uniformly across single cells. **a** Number of unique (blue) and shared (yellow) genes upregulated after stimulation (6 h, 10 ng/ml LPS+10 ng/ml IFN-γ or 100 ng/ml IL-4). **b** Volcano plot of differential gene expression after co-stimulation (6 h, 10 ng/ml LPS+10 ng/ml IFN-γ and 100 ng/ml IL-4) relative to LPS+IFN-γ alone (left) and IL-4 alone (right). Dark blue dots (see key) indicate the unique core genes (UCGs) selectively induced by LPS+IFN-γ or IL-4, respectively. Cyan dots indicate shared genes induced by both LPS+IFN-γ or IL-4. Yellow dots indicate UCGs modulated by co-stimulation, identified by an FDR ≤ 0.05 and change in expression ? 1.5-fold relative to expression after stimulation with LPS+IFN-γ or IL-4 alone. FDR determined as in Fig. 1. **c** Transcript levels for indicated targets in BMDMs after stimulation for 6 h with media alone, LPS+IFN-γ, IL-4, or both, measured by population RT-qPCR (left) and scRNA-seq (right). Population mRNA levels are presented relative to those of the control gene *Ubc* (mean ± SEM, n=3). Unpaired *t*-test; *\*p* ≤ 0.05, *\*\*\*p* ≤ 0.001, n.s. = not significant. Single-cell data is presented as the ln(transcript count +1) from a single experiment. **d** Noise in gene expression (calculated by Fano factor) across single cells stimulated with LPS+IFN-γ versus co-stimulated cells (left) or IL-4 vs co-stimulated cells (right) for genes negatively (blue) and positively (red) regulated by co-stimulation. **e** Change in cell-to-cell gene expression noise of LPS+IFN-γ-induced genes (left) and IL-4-induced genes (right) in co-stimulated cells relative to single stimulation.

We compared observations from our single-cell dataset to population-level RT-qPCR data on co-stimulated cells for canonical LPS+IFN-γ and IL-4-induced core genes. Population-level measurements confirmed our scRNA-seq results, with LPS+IFN-γ-induced genes *Il12b* and *Il6* and IL-4-induced genes *Arg1* and *Chil3* exhibiting sensitivity to cross-inhibition, while other LPS+IFN-γ-induced genes, including *Nos2* and *Tnf* were not sensitive to co-stimulation (**Fig. 2c**, bar plots). Inhibition or resistance to co-stimulation for this set of target genes was also observed at the protein level at 6 and 24 hours (**Supplementary Fig. 2a-b**).

Importantly, although we observed substantial cross-inhibition in the cell population for some gene targets, the scRNA-seq data revealed that there was substantial cell-to-cell heterogeneity, such that some individual cells exhibited transcript levels that appeared uninhibited even after stimulation with both cues (**Fig. 2c**, violin plots). Given the significant cell-to-cell heterogeneity in cross-inhibition of the gene programs, one hypothesis is that macrophages are diversifying along these negatively regulated pathways. To identify cross-inhibited genes that are most variably expressed, we calculated the noise in gene expression as measured by Fano factor (variance divided by mean) for the LPS+IFN-γ- and IL-4-induced genes in both single and co-stimulated cells. Interestingly, we observed a subset of inhibited genes that upon co-stimulation decrease their mean expression across the population while increasing gene expression noise (**Fig. 2d** and **Supplementary Fig. 2c**). Strikingly, 9 out of the top 15 genes induced by LPS+IFN-γ and exhibiting increased noise were secreted cytokines or chemokines, including the T-cell polarizing inflammatory cytokines *Il6* and *Il12b* (**Fig. 2e**, left). Notably, this gene set also includes NF-κB inhibitor zeta *(Nfkbiz),* which is known to stimulate the transcription of a subset of inflammatory response genes including *Il6* and *Il12b*^27^. IL-4-induced genes exhibiting increased noise included canonical genes like *Arg1* and *Socs2,* as well as transcription factors associated with promoting anti-inflammatory phenotypes like *Mafb* and *Klf4* (**Fig. 2e**, right)^28,29^. Altogether, these results suggest extensive transcriptional cross-regulation between the LPS+IFN-γ and IL-4 core gene programs that are gene-specific and that vary substantially from cell to cell.

We next asked if cross-inhibited genes were enriched for pathways regulating certain cellular functions. The set of cross-inhibited genes from each single-stimulus program was analyzed using Ingenuity Pathway Analysis (IPA) to determine what cell functions are most sensitive to inhibition by co-stimulation (**Supplementary Fig. 2d**). LPS+IFN-γ-induced genes that were inhibited by co-stimulation were significantly enriched for pathways involved in communication between innate and adaptive immune cells and regulation of cytokine production, consistent with the targets identified in the noise analysis. IL-4-induced genes subject to inhibition by co-stimulation were enriched for STAT3 and JAK2 in hormone-like cytokine signaling, although the results were less significant than for LPS+IFN-γ. Altogether, the IPA result suggested that important functions typically stimulated by LPS+IFN-γ or IL-4, including secretion of cytokines and chemokines and STAT3 signaling, are inhibited by co-stimulation.

### Machine learning classification suggests that LPS+IFN-γ-induced and IL-4-induced gene programs subject to negative cross inhibition are selectively expressed in single cells after co-stimulation

The extensive cell-to-cell heterogeneity and negative correlations between LPS+IFN-γ and IL-4 observed in single cells suggest that some single cells might selectively express sets of LPS+IFN-γ-induced or IL-4-induced genes despite having been stimulated by both cues. To explore this hypothesis, we used a neural network (NN) to classify our co-stimulated cells into LPS+IFN-γ, IL-4, or mixed cell states after training a classifier on different sets of genes chosen to define the LPS+IFN-γ and IL-4 transcriptional response signatures. Specifically, we used cells stimulated with LPS+IFN-γ alone and IL-4 alone to train two separate classifiers to identify these single-stimulation signatures (**Fig. 3a**). We then used these classifiers to predict the cell states of our co-stimulated cells, which could be classified as LPS+IFN-γ-dominant, IL-4-dominant, Mixed (i.e., both gene programs are detected), or Unclassified. We first trained the NN classifiers using the full transcriptome, the top 200 unique core genes (UCGs), or the top 50 UCGs. When using both the global transcriptome or different numbers of UCGs, the NN predicted that most cells were dominated by the LPS+IFN-γ gene program, with just a few cells exhibiting a mixed state and no cells exhibiting an IL-4-dominant signature (**Fig. 3b**, left). However, when we restricted our training set to UCGs downregulated with co-stimulation, the number of co-stimulated cells classified as exhibiting an LPS+IFN-γ-dominant state was reduced and a fraction of cells exhibiting an IL-4-dominant state emerged (**Fig. 3b**, right). Interestingly, the fractions of co-stimulated cells classified as LPS+IFN-γ-dominant or IL-4-dominant were both substantially larger than the Mixed and Unclassified fractions. This result suggests that a subset of co-stimulated macrophages behave as if responding to only one stimulus for negatively cross-regulated transcriptional programs.

**Fig. 3.**
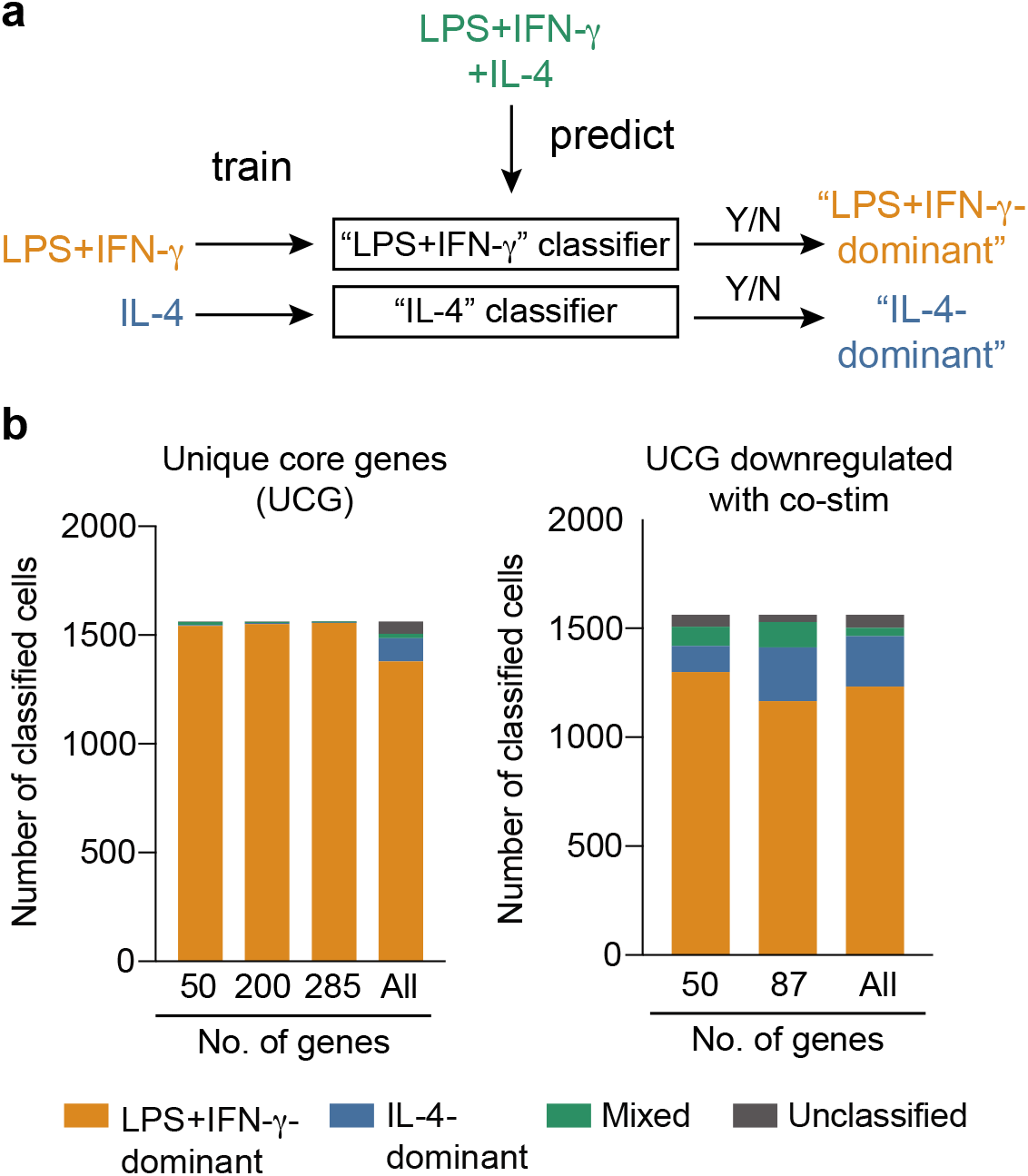
Neural network classifier suggests cross-inhibited genes are selectively expressed by single cells after co-stimulation. **a** Schematic of the neural network classifiers built to categorize cells as LPS+IFN-γ-dominant, IL-4-domi-nant or Mixed. **b** Results of the neural network classifier in identifying the transcriptional state of co-stimulated cells. Bars represent the number of cells identified as LPS+IFN-γ-dominant (yellow); IL-4-dominant (blue); Mixed (green); or Unclassified (grey). The classifiers were trained on either unique core genes (UCGs, left), or UCGs that displayed cross-inhibition after co-stimulation (right).

### *Il6* and *Il12b* and their transcription factor *Nfkbiz* are expressed orthogonally with *Arg1* and Chil3 and their transcription factor *Klf4*

Negative cross-regulation has the potential to create specialized functions within a subset of macrophages. If cross-regulation between two genes is strong, then these genes would not be anti-correlated but rather would be orthogonally expressed (i.e., the presence of gene A would significantly reduce the probability of expressing gene B such that they would not be expressed together). This expression pattern may be exacerbated in scRNA-seq data for genes with low transcript numbers due to dropout^30^. To quantify orthogonal expression, we expressed the scRNA-seq data as binary data with a threshold of 2 detected transcripts (see **Methods**) and calculated the odds-ratio between all pair-wise combinations of down-regulated genes. The odds ratio is the ratio of the odds of expressing A in the presence of B to the odds of expressing A in the absence of B. If the log of the odds ratio is negative, then the presence of B decreases the likelihood of A and is more likely to exhibit orthogonal gene expression.

We found that genes exhibiting significant negative odds ratios were relatively few, and were more likely to be observed between core genes upregulated by LPS+IFN-γ versus IL-4 than between core genes of the same program (**Fig. 4a**). We noted that transcripts for the T cell-polarizing cytokines *Il6* and *Il12b* had negative odds ratios with *Arg1* (**Fig. 4a**). To confirm orthogonal expression of these two pathways, we plotted the sc-RNAseq expression for *Il6* and *Il12b* against *Arg1* across all conditions (**Fig. 4b**). Plotting single-cell expression of these transcripts revealed a striking orthogonal expression pattern: cells with high levels of *Il6* or *Il12b* had no or very low expression of *Arg1,* and vice versa (**Fig. 4b**). Orthogonal expression with *Il12b* and *Il6* was also observed with the IL-4-associated target *Chil3* (**Supplementary Fig. 3a**). Thus, it appears that select LPS+IFN-γ and IL-4 targets are rarely expressed coincidentally in the same individual cell.

**Fig. 4.**
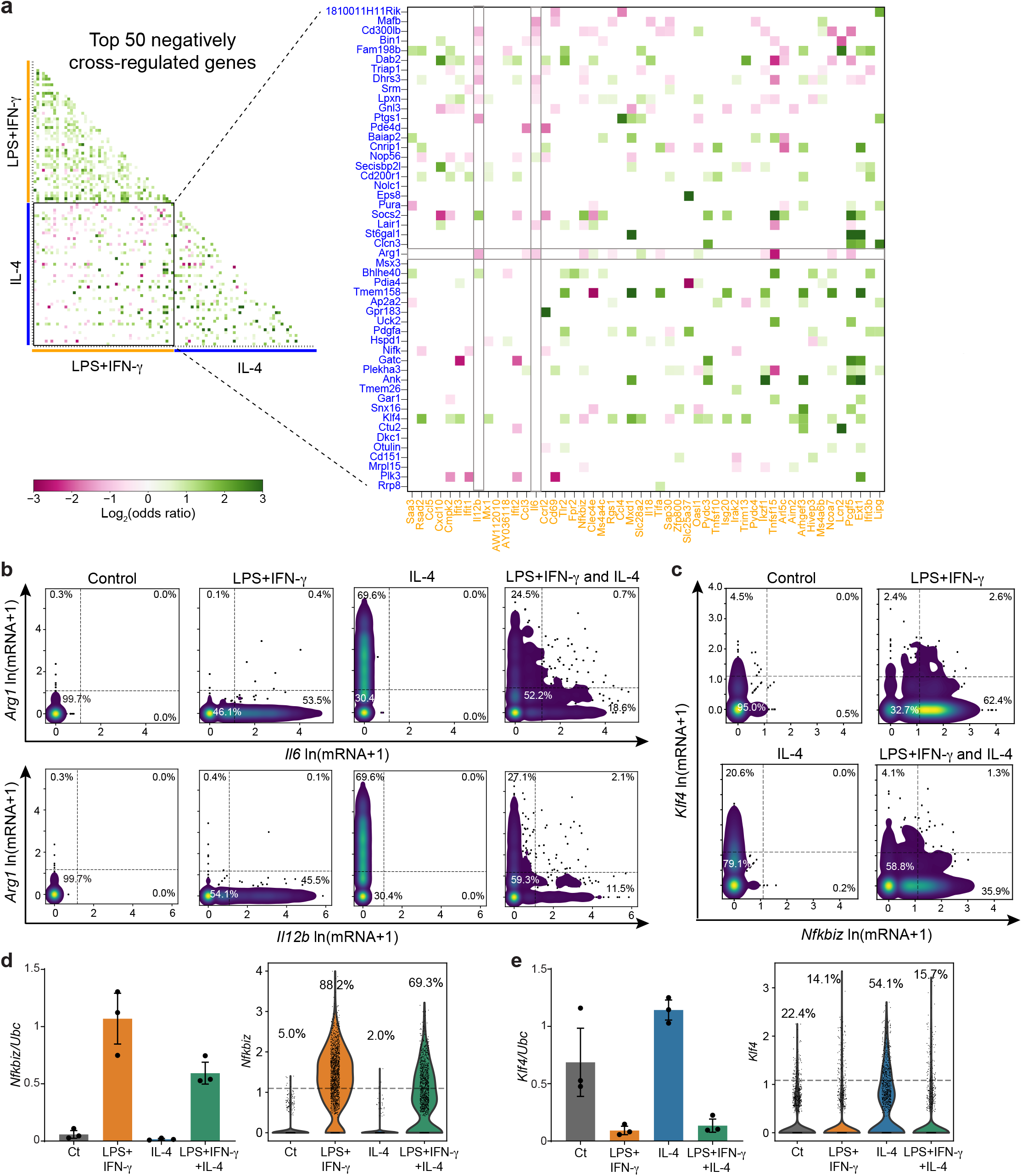
*Il6* and *Il12b* and their transcription factor *Nfkbiz* are expressed orthogonally with *Arg1* and its transcription factor *Klf4.* **a** Pairwise odds ratios for the top 50 downregulated genes in the LPS+IFN-γ (yellow) and IL-4 (blue) core gene programs measured in co-stimulated cells. Data are presented as the log_2_(odds ratio). Color bar indicates the strength and magnitude of the association. Odds ratios with a *p*-value > 0.05 (determined by Fisher’s exact test) are set to 0. **b-c** Density scatter plots of scRNA-seq transcript counts across individual cells for *Arg1* vs. *Il6* (**b**; top) and *Il12b* (**b**; bottom) and *Klf4* vs. *Nfkbiz* (**c**). Data is represented as ln(transcript count +1). **d-e** Population RT-qPCR measurements (left) and violin plots of scRNA-seq measurements (right) of *Nfkbiz* (**d**) and *Klf4* (**e**) after stimulation for 6 h with media alone, LPS+IFN-γ, IL-4, or both. Population mRNA levels are presented relative to those of the control gene *Ubc* (mean ± SEM, n=3). Single-cell data is presented as the ln(transcript count +1) from a single experiment.

We next looked for transcription factors related to these targets that might play a role in selective expression. We observed that *Nfkbiz,* a positive transcription factor for the secondary response genes *Il6* and *Il12b*^19^, also had a negative odds ratio with *Arg1,* suggesting that this pathway may be orthogonally regulated with the *Arg1* pathway (**Fig. 4a**). In support of this hypothesis, *Nfkbiz* had a negative odds ratio with *Klf4,* a positive transcription factor for *Arg1* and *Chil3* that also negatively regulates LPS-stimulated pro-inflammatory genes (**Fig. 4a, c**)^29^. Importantly, we previously noted that *Nfkbiz* and *Klf4* are transcription factors that are subject to negative cross-regulation and increase their noise after co-stimulation (**Fig. 2e**). We then measured the expression of these transcription factors by RT-qPCR in a cell population and confirmed that *Nfkbiz* and *Klf4* are inhibited after co-stimulation (**Fig. 4d-e**). Together these results suggest that macrophages co-stimulated with LPS+IFN-γ and IL-4 may orthogonally express the transcription factors *Nfkbiz* and *Klf4,* resulting in orthogonal expression of *Arg1* and *Chil3* versus the secondary cytokine genes *Il6* and *Il12b.*

We did not observe orthogonal expression with *Arg1* or *Klf4* for transcripts encoding primary cytokines and chemokines including *Ccl5* and *Tnf* (**Supplementary Fig. 3b**). This is consistent with the fact that expression of *Tnf* and other primary genes is *Nfkbiz*-independent^27,31^. Overall, our results strongly suggest that co-stimulated macrophages specialize in the production of transcripts for *Arg1* and *Chil3* versus *Il6* and *Il12b*, but not primary secreted cytokines/chemokines, at least partially through the heterogeneous regulation of *Klf4* and *Nfkbiz* expression.

### Orthogonal gene expression in co-stimulated cells leads to macrophage subsets secreting either IL-6 and IL-12p40 or Chi3l3 at 48 hours

Although our results demonstrate orthogonal expression of *Klf4* and *Nfkbiz* transcripts and their targets *Arg1* and *Chil3* or *Il6* and *Il12b,* respectively, in LPS+IFN-γ and IL-4 co-stimulated macrophages at 6 hours, it is unclear if this specialization is sustained long enough to direct macrophages to have distinct functions. To explore this possibility, we used a multiplexed singlecell secretion assay that we previously used to measure the heterogeneity of LPS-stimulated macrophages^16,24^. We stimulated BMDMs cultured in the single-cell device with LPS+IFN-γ, IL-4, or a combination of LPS+IFN-γ+IL-4, and captured the secretion of 11 different proteins for 48 hours to obtain an integrative measurement of the final secretion state for each macrophage. We measured a combination of inflammatory secreted proteins including TNFα, CCL5, IL-6, and IL-12p40. Because Arg1 is not secreted, we measured Chi3l3 (the protein product of *Chil3*) as the primary secreted protein in response to IL-4 stimulation.

After 48 hours of co-stimulation, we observed orthogonal secretion of IL-6 and IL-12p40 and the IL-4-induced marker Chi3l3 (**Fig. 5a**), similar to what we observed at the transcript level at 6 hours. Indeed, we found very few cells that co-secreted IL-6 and Chi3l3 or IL-12p40 and Chi3l3 (**Fig. 5b**, green). This orthogonal behavior was not observed between CCL5 and Chi3l3 or TNFα and Chi3l3 (**Supplementary Fig. 4a**), consistent with the scRNA-seq results (**Supplementary Fig. 4b**). To verify orthogonal secretion, we calculated the odds-ratio between all pair-wise combinations of secreted proteins in co-stimulated cells to measure whether the secretion of a given protein is more, less, or as likely in a single cell when that cell is secreting the paired protein. We observed that the only pair of proteins with a negative log_2_(odds ratio) was IL-6 and Chi3l3 (odds ratio = 0.44 and log_2_(odds ratio) = −1.2), indicating that if a cell is secreting IL-6, the odds of that cell also secreting Chi3l3 are 56% lower than the odds of secreting Chi3l3 when IL-6 is not being secreted (**Fig. 5c**). These data demonstrate that the specialization observed in transcription at 6 hours is maintained for secretion at 48 hours, such that co-stimulated cells either secrete IL-6 and IL-12p40 or Chi3l3.

**Fig. 5.**
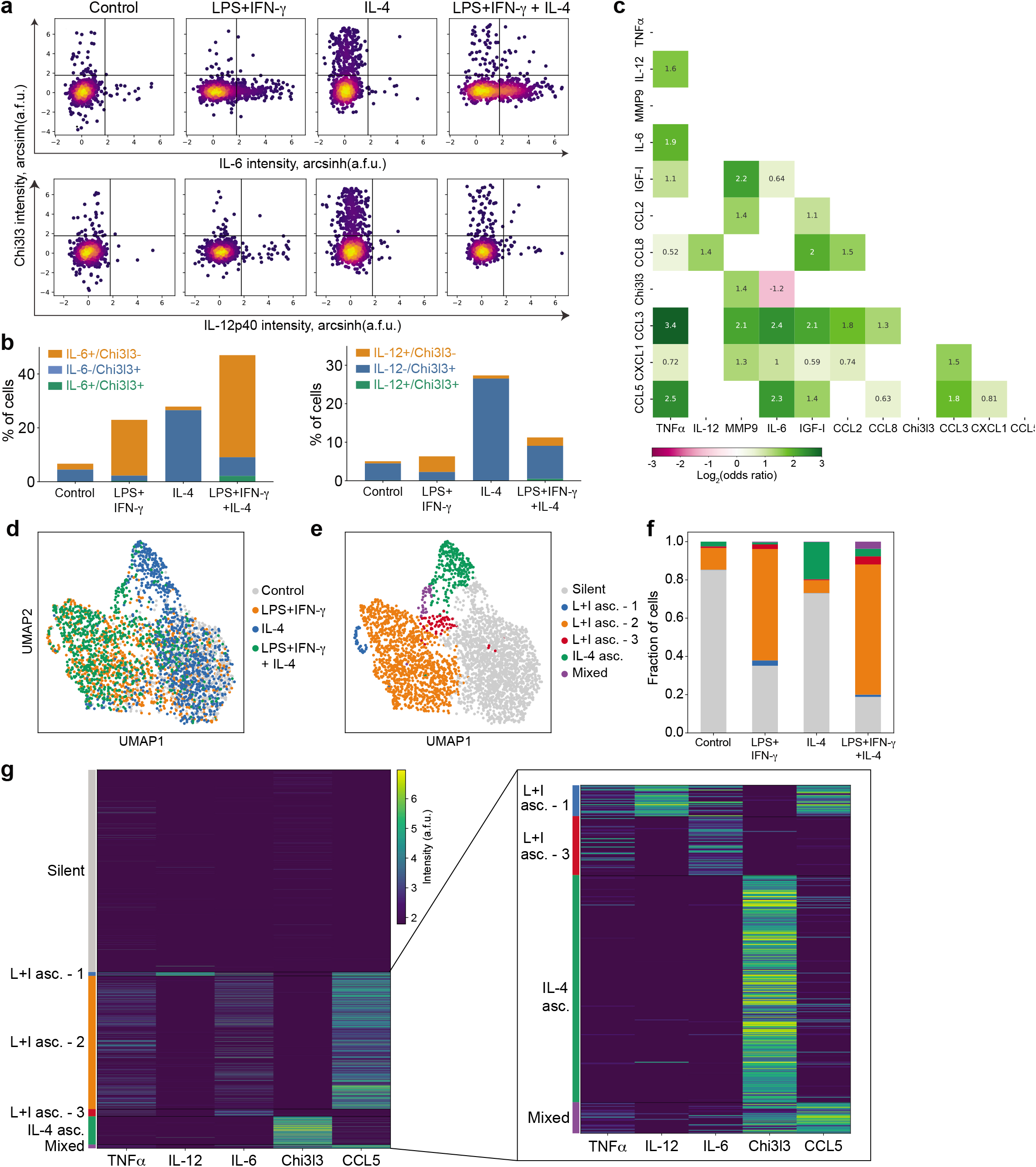
Single-cell secretion assay identifies long-term specialization in co-stimulated cells. A multiplexed single-cell secretion assay was used to measure cytokine/chemokine production in individual BMDMs stimulated for 48 h with media alone, 10 ng/ml LPS+10 ng/ml IFN-γ, 100 ng/ml IL-4, or both. **a** Density scatter plots of single-cell secretion intensity (a.f.u.) across individual cells for Chi3l3 vs IL-6 (top) and Chi3l3 vs IL-12p40 (bottom). **b** Quantification of single- and double-positive cells after co-stimulation for Chi3l3 and IL-12p40 (left) and Chi3l3 and IL-6 (right). **c** Pairwise odds ratios for all secreted proteins measured in co-stimulated cells. Data are presented as the log_2_(odds ratio). Color bar indicates the direction and magnitude of the association. Odds ratios with an associated *p*-value > 0.05 (determined by Fisher’s exact test) are set to 0. **d-e** UMAP visualization of single-cell secretion data for 5 proteins (Chi3l3, CCL5, IL-6, IL-12p40, and TNFα), colored by stimulation (**d**) or consensus cluster (**e**). **f** Quantification of the fraction of cells present in each consensus cluster identified by ensemble clustering. **g** Heatmap of secretion profiles for the five secreted proteins in (**d**) grouped by consensus cluster. Inset is a magnified version of the smaller clusters. All single-cell secretion data presented is pooled from 2 biological replicates. L+I asc. = LPS+IFN-γ-associated; IL-4 asc. = IL-4-associated.

Finally, we explored the extent to which subsets of co-stimulated macrophages showed secretion programs skewed towards LPS+IFN-γ versus IL-4 stimulation. To test this, we performed dimensionality reduction and unsupervised ensemble clustering^32^ on five proteins distinguishing LPS+IFN-γ- and IL-4-stimulated cells in the single-cell secretion dataset (specifically: TNFα, CCL5, IL-6, IL-12p40, and Chi3l3). We found that the cells stimulated with individual stimuli divided into two separate branches in the low-dimensional UMAP projection, while the co-stimulated cells spread across these two branches (**Fig. 5d**). Clustering analysis revealed that all conditions contained a cluster of silent cells with low secretion of all proteins (**Fig. 5e-g**). Additionally, cells stimulated with LPS+IFN-γ contained clusters of cells secreting a combination of TNFα, IL-12p40, IL-6 and CCL5 (**Fig. 5e-g**). Cells stimulated with IL-4 were composed of mostly silent cells, as well as a distinct cluster secreting only Chi3l3 (**Fig. 5e-g**). Co-stimulated cells were composed of a distribution of the three LPS+IFN-γ-associated clusters and the IL-4-associated cluster, as well as a small Mixed cluster with cells co-secreting Chi3l3, CCL5, and TNFα (**Fig. 5e-g**). Overall, we conclude that for secretion, a majority of macrophages co-stimulated with LPS+IFN-γ and IL-4 exhibit secretion profiles consistent with only one of these stimuli.

## Discussion

Macrophages exhibit diverse polarization states *in vivo* in response to complex cues in their microenvironments, but our understanding of how individual macrophages respond to simultaneous cues is limited. In this study, we used a combination of population and single-cell measurements and computational analyses to explore how macrophages respond when simultaneously presented with LPS+IFN-γ and IL-4. We found that while co-stimulated macrophages displayed a distinct global transcriptional program, variable negative crossregulation between some LPS+IFN-γ- and IL-4-stimulated gene programs resulted in significant cell-to-cell variability, such that some co-stimulated macrophages are skewed towards one of the two transcriptional programs. In particular, our results suggest that negative cross-regulation by the transcription factors *Klf4* and *Nfkbiz* leads to the orthogonal expression of the secondary Th1 cytokines *Il6* and *Il12b* with *Arg1* and *Chil3,* and that this results in macrophages with specialized secretion functions that are maintained for at least 48 hours.

The cross-regulation observed between LPS+IFN-γ and IL-4 programs was observed at various time-points, concentrations, and at both transcription and secretion levels. The direction and magnitude of cross-regulation was gene-dependent and observed at the single-cell level as well as in macrophage populations. Interestingly, co-stimulation induced substantial cell-to-cell heterogeneity in the transcriptional response, such that some cells display reduced transcript levels of certain genes, while other cells remain completely uninhibited. This heterogeneity suggests a diversification in the population when presented with different cues, and could be a viable strategy to deal with complex microenvironments.

To further explore this diversification, we used a combination of correlation analysis, neural network classifiers and other statistical analyses to examine the cell identity of co-stimulated cells. We found that the polarization state identified by these analyses was dependent on the transcriptional program analyzed. When looking at a global transcriptional state, most cells are classified as mixed cells; however, when we looked at genes induced by LPS+IFN-γ that are susceptible to cross-inhibition, a subset of co-stimulated cells appeared more specialized. Indeed, we show that some of the most widely accepted markers for M1-like and M2-like polarization states can classify co-stimulated cells as specialized, whereas others would classify as mixed. Thus, in agreement with previous reports, our results argue that overall polarization is better understood as a spectrum; however, our results add the novel insight that negative cross-regulation significantly reduces the probability that individual macrophages exhibit an intermediate state in this spectrum for a subset of gene programs^3,4^. This gene set-dependent classification may partially explain the difficulty in assigning polarization states *in vivo* – indeed, many studies find contradicting results when trying to identify the polarization state of many tissue-resident macrophages^33,34^.

Our observation that the transcription factors *Klf4* and *Nfkbiz* are orthogonally expressed suggests a regulatory mechanism for specialization of IL-6 and IL-12p40 secretion versus Chi3l3 secretion. *Klf4* is an example of a macrophage-specific transcription factor that regulates specialized gene programs^35^. Based on sequence analysis, there is a binding site for *Klf4* on the promoter of *Nfkbiz^35^,* and thus it is possible that *Klf4* directly negatively regulates *Nfkbiz* upon co-stimulation. Recent work by Piccolo et al. identified several transcriptional and epigenetic mechanisms that regulate the integration of IFN-γ and IL-4 signals and are responsible for their cross-regulation^14^. Furthermore, recent developments in single-cell epigenetic measurements have identified how cell-to-cell variation can be encoded via chromatin accessibility and can direct complex biological decisions like hematopoietic differentiation^33,34^. Future work can focus on measuring differences in the accessibility of distinct promoters, which could explain the observed specialization and help answer if these partially specialized cells exist in a predisposed state before stimulation, or if the specialization is imposed and reinforced after receiving the stimuli.

Macrophage-secreted cytokines and chemokines are essential for coordinating the immune response in complex tissue microenvironments. Interestingly, we found that LPS+IFN-γ-stimulated genes that were negatively regulated by IL-4 were enriched for secreted cytokines and chemokines and this was associated with increased gene expression noise (**Fig. 2e** and **Supplementary Fig. 2d**). Importantly, we observed nearly orthogonal secretion of IL-4-stimulated Chi3l3 and LPS+IFN-γ-stimulated IL-6 and IL-12p40 (**Fig. 5a**), which are essential cytokines for inducing T cell polarization^35^. The observed diversification of single macrophages across a range of secretion programs when presented with opposing cues may enable macrophages to more quickly adapt to changing environments. This concept of ‘bet-hedging’ has been described in many multi-cellular systems as a way to increase the robustness of the population, including bacterial responses to resource availability as well as mammalian diversification of NF-kB signaling or T cell activation^36–38^. It is possible that macrophages display this partial specialization as a way to execute the dynamic nature of immune responses, such as the quick transition observed during the resolution of inflammation, where macrophages have to coordinate a successful change from an anti-microbial to a tissue-repair phenotype. Indeed, the ability of macrophages to quickly adapt to changing environments has been exploited to design therapies that induce anti-tumor immunity in melanoma^19^.

The single-cell response of BMDMs to co-stimulation with LPS+IFN-γ and IL-4 has important implications for understanding macrophage responses to complex cues that co-exist *in vivo.* Overall our results suggest that variability in the intracellular networks that negatively crossregulate polarization programs provides an important source of intercellular heterogeneity among macrophages. However, the importance of these intracellular networks relative to the various contributors to heterogeneity *in vivo* is not yet clear. Due to the limited diffusion distances and communication capacity of *in vivo* settings, it has become apparent that macrophage heterogeneity in the body is at least partially explained by the local microenvironment around individual macrophages^16,17^. For example, different locations within tumor microenvironments can induce functionally distinct macrophages that are important in regulating disease progression^19,39^. It is therefore possible that *in vivo*, some of the observed heterogeneity and specialization is location dependent. In addition to genetic and local regulation, recent work has highlighted how metabolic state and circadian rhythm have profound effects on macrophage functions^40–42^. It is possible that differences in metabolic states and molecular clocks help generate the diversity required to induce partially specialized macrophages after co-stimulation. Thus, determining the relative importance of non-genetic cell-to-cell heterogeneity vs. other in vivo factors will be an important future direction.

## Materials and Methods

### Mice and cell culture

Wild type C57BL/6J mice purchased from Jackson Laboratories were used. Bone marrow-derived macrophages were generated as previously described^43^. Briefly, bone marrow was extracted from the hind legs of the mouse with a syringe. After red blood cell lysis with ammonium-chloride-potassium lysis buffer (Lonza), cells were incubated for 4 hours at 37° C with 5% CO2 in a nontissue culture (TC) treated plastic petri dish with BMDM media (RPMI supplemented with 10% FBS, 100 U/ml penicillin, 100 μg/ml streptomycin, 1% sodium pyruvate, 25 mM HEPES buffer, 2 mM L-glutamine, and 50 μM 2-mercaptoethanol). After 4 hours, the non-adherent cells were transferred to a new petri dish and incubated with BMDM media + 20 ng/ml macrophage-colony stimulating factor (M-CSF; Peprotech). After 3 days, an additional 10 ml of BMDM media + 20 ng/ml M-CSF was added to the plate. 6 days after plating, cells were harvested in PBS + 5 mM EDTA with gentle scraping, and the cell suspension was used to seed new non-TC treated dishes or microwell devices. All mice were housed in the Yale Animal Resources Center in specific pathogen-free conditions. All animal experiments were performed according to the approved protocols of the Yale University Institutional Animal Care and Use Committee.

### *In vitro* BMDM experiments

BMDMs were plated and stimulated in BMDM media + 10 ng/ml M-CSF. Cells were plated in non-TC treated 6- or 12-well plates (Falcon) at a density of 100,000 cells per cm^2^ and allowed to adhere overnight. Cells were then stimulated at the indicated doses and times using LPS (Invivogen), IFN-γ (Peprotech) and/or IL-4 (Peprotech).

### Quantification of secretion in population

To measure secretion in populations of cells, supernatant from stimulated BMDMs was collected and stored at 4°C for no more than a week. Secreted protein levels were measured using enzyme-linked immunosorbent assay (ELISA) kits according to the manufacturer’s recommendations. Each ELISA kit reference number and manufacturer are listed in **Table 1**.

**Table 1.** List of capture and detection antibody pairs for ELISA secretion profiling.

| Antibody Pair | Vendor | Catalog No. |
| --- | --- | --- |
| TNFα | eBioscience | 88-7324-88 |
| CCL17 | R&D | DY529 |
| IL-12p40 | BD Biosciences | 555165 |
| IL-10 | BD Biosciences | 555252 |
| MMP9 | R&D | DY6718 |
| IL-6 | R&D | DY406 |
| IGF-I | R&D | DY791 |
| CCL2 | R&D | DY479 |
| CCL8 | R&D | DY790 |
| Chi3l3 | R&D | DY2446 |
| CCL3 | R&D | DY450 |
| CXCL1 | R&D | DY453 |
| GM-CSF | R&D | DY415 |
| CCL22 | R&D | DY439 |
| CCL5 | R&D | DY478 |

### RT-qPCR

RT-qPCR was performed as previously described^44^. Briefly, RNA was extracted using the RNEasy Mini Kit (Qiagen). Genomic DNA was removed on-column with RNase-free DNase (Qiagen) or with the TURBO DNA-free kit (Ambion), and complementary DNA (cDNA) was synthesized using a dT oligo primer and Superscript III RT (Invitrogen). After dilution in nuclease-free water, cDNA was quantified using SYBR-green for quantitative reverse transcription polymerase chain reaction on a CFX Connect Real-Time System (Bio-Rad) with the following amplification scheme: 95°C denaturation for 1.5 minutes followed by 40 cycles of 95°C denaturation for 10 secs, 65°C annealing for 10 secs, and 72°C elongation for 45 secs with a fluorescence read at the end of the elongation step. This was followed by a 65°C to 85°C melt-curve analysis with 0.5°C increments. All samples were normalized to house-keeping gene *Ubc* (ubiquitin). Primers are reported in **Table 2**.

**Table 2.**
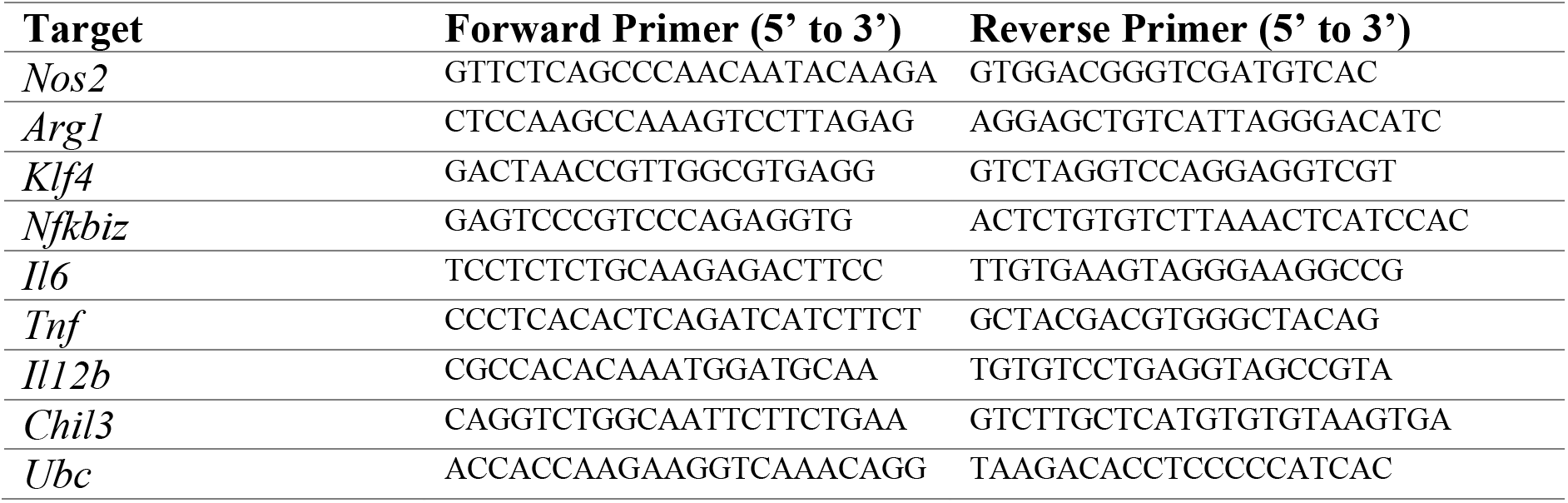
List of primers used for qPCR.

### Flow Cytometry

Macrophages were lifted with gentle scraping in ice-cold PBS + 5 mM EDTA. For intracellular protein staining, cells were blocked with Fc receptor (CD16/CD32) antibody (eBioscience, clone 93) on ice for 15 mins in FACS buffer (PBS + 2% FBS). Then the cells were fixed with Cytofix/Cytoperm (BD Biosciences) and stained in 50 μl with anti-Nos2-AlexaFluor488 at 1:500 dilution (eBioscience, clone CXNFT) and anti-Arg1-APC at 1:10 dilution (R&D, polyclonal) for an hour at 4°C. For phospho-flow, cells were fixed immediately after lifting with PhosFlow Fix buffer for 10 mins at 37°C, and subsequently permeabilized with PhosFlow Perm Buffer III for thirty minutes on ice (BD Biosciences). Cell suspensions were then blocked with Fc receptor antibody as above, and stained with anti-pSTAT1(Y701)-AlexFluor488 (Cell Signaling Technologies, clone 58D6) and anti-pSTAT6(Y641)-AlexaFluor647 (Cell Signaling Technologies, clone D8S9Y). All data was acquired on an Accuri B6 flow cytometer (BD Biosciences), and analyzed with FlowJo (FlowJO, LLC).

### Single-cell RNA sequencing

Stimulated BMDMs were lifted in ice-cold PBS+ 5 mM EDTA with gentle scraping, washed, counted, and immediately processed at low density for scRNA-seq using the 10X platform (10X Genomics). Library construction and sequencing was performed by the Yale Center for Genomic Analysis according to the manufacturer’s recommendations.

### scRNA-seq analysis

The sequencing data was processed using the standard cellranger pipeline (10X Genomics). Further downstream analysis was performed using the Python package scanpy^45^. Cells were filtered for quality control to avoid doublets and dead cells, and counts were normalized using the scran package in R in a standard processing pipeline previously described^46,47^. Dimensionality reduction and visualization was performed within the scanpy package. Correlation and odds-ratio analysis were performed using custom scripts in Python. To find core genes for each polarization program, differential testing using Wilcoxon rank-sum test with Benjamini-Hochberg correction for multiple comparisons was performed to find differentially expressed genes between stimulated cells and control cells. Core genes were defined as genes with a minimum fold change of 1.5, maximum FDR of 0.05, and expressed in at least 15% of the stimulated cells. From this list of core genes, we found the UCGs that were only induced by a single stimulation, and not by both.

### Microwell assay for single-cell secretion profiling

The single-cell secretion profiling experiments were performed as previously described^10,22^, with some modifications for the analysis of primary mouse BMDMs. In brief, the capture antibodies (Table 3.2) were flow patterned onto epoxysilane-coated glass slides (SuperChip; ThermoFisher). The polydimethylsiloxane nanowell arrays and antibody barcode glass slides were blocked using complete BMDM media. Fully differentiated BMDMs were resuspended in complete BMDM media and 10 ng/ml M-CSF and supplemented with 125 nM of live cell marker (Calcein AM; ThermoFisher) to facilitate automatic live cell detection. The cells were added to the device and allowed to adhere overnight. The next day, cells were stimulated and covered with the antibody barcode slide, secured with screws, and allowed to incubate for 48 hours. At the end of the incubation period, the device was imaged with an automated inverted microscope (Eclipse Ti; Nikon or Axio Observer Z1; Zeiss) to record well position and cell locations. The device was then disassembled to perform the sandwich immunoassay. The glass slide was incubated with a mixture of detection antibodies (Table 3.2) for 1 h, followed by an incubation with 20 μg/ml streptavidin-APC (eBioscience) for 30 min, rinsed with PBS and water, and finally scanned with a Genepix 4200A scanner (Molecular Devices).

### Single-cell secretion profiling data processing

Device images were analyzed using a custom script in MATLAB (MathWorks) to automatically detect well location and number of cells per well, extract all signals from each well, and process the data (https://github.com/Miller-JensenLab/Single-Cell-Analysis). In brief, after automatic well and live cell detection, signal image registration, and manual curation, the software automatically extracted the intensity signal from each antibody for all the nanowells in the device. The signal across the chip for each individual antibody was normalized by subtracting a moving Gaussian curve fitted to the local zero-cell well intensity levels. A secretion threshold for each antibody was then set at the 99th percentile of all normalized zero-cell wells. Finally, the data were transformed using the inverse hyperbolic sine with a cofactor set at 0.8× secretion threshold. To further visualize the data, custom Python scripts were used to generate UMAP visualizations and density scatter plots. Odds ratios were calculated using the same methods as for scRNA-seq data.

### Neural network classifier

The neural network classifier was built using the machine learning python package scikit-learn^48^. The data from the cells receiving a single stimulus was split into a training and a testing dataset to build and test the classifier. A one-vs-the-rest (OvR) multilabel classifier strategy was used to enable non-mutually exclusive labels and identify mixed cells. Briefly, two multi-layer perceptron (MLP) classifiers were trained on the LPS+IFN-γ- and IL-4-stimulated cells to distinguish cells stimulated with each cue. The MLPs had three hidden layers and used the hyperbolic tan function as its activation function. Then the OvR multilabel classifier was used to predict the identity of co-stimulated cells, which labelled each cell as LPS+IFN-γ-stimulated, IL-4-stimulated, both, or none (no prediction).

### Ensemble clustering

Ensemble clustering was performed with a python package from the Naegle lab (https://github.com/NaegleLab/OpenEnsembles)^32,49^. Briefly, clustering was performed by sweeping across multiple clustering algorithms (Affinity Propagation, agglomerative, spectral, Birch, kmeans), across several distance and linkage metrics (average, complete, euclidean, cosine, ward, l1 and l2) and across a wide range of k values (that determines the number of clusters to identify). A consensus clustering solution was found by linking the co-occurrence matrix^32,49^. Clusters with less than 3 cells were discarded for analysis.

### Statistics

Data were presented as mean ± SEM unless otherwise specified. Statistical analysis was performed by Student’s t test or one-way ANOVA and the Sidak method of correction for pairwise multiple comparisons as specified in the figure legends. Normal and equal distribution of variances was assumed. Values were considered significant at *P* < 0.05 and are indicated as *, *P* < 0.05; **, *P* < 0.01; ***; *P* < 0.001; and ****, *P* < 0.0001. All analyses were performed using Prism version 7.0 software (GraphPad) or custom python scripts.

## Acknowledgements

The authors thank Ruslan Medzhitov and Andre Levchenko for their insightful comments and discussion, as well as all members of the Miller-Jensen lab for helpful discussion and experimental advice. This work was supported by the National Institutes of Health (R01-GM123011 to K.M-J.).

## Author contributions

Conception: A.R. Muñoz-Rojas, and K. Miller-Jensen. Development of methodology: A.R. Muñoz-Rojas and K. Miller-Jensen. Acquisition of data: A.R. Muñoz-Rojas, I. Kelsey, and J. Pappalardo. Analysis and interpretation of data: A.R. Muñoz-Rojas, I. Kelsey, and K. Miller-Jensen. Writing, review, and/or revision of the manuscript: A.R. Muñoz-Rojas, I. Kelsey, and K. Miller-Jensen. Study supervision: K. Miller-Jensen.

---

**Supplementary Fig. 1 (associated with Fig. 1).**
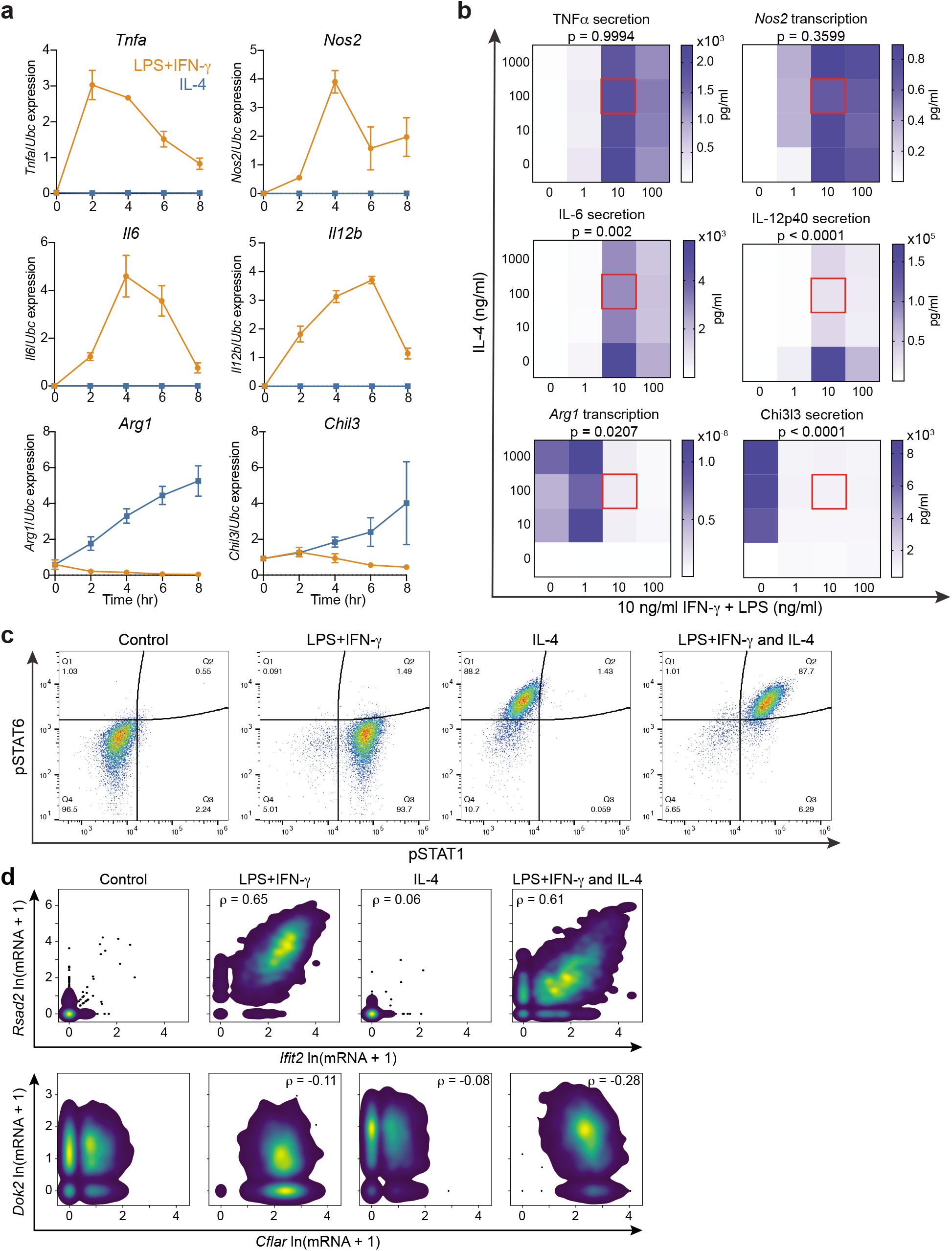
Timing and dose dependence of cross-inhibition between LPS+IFN-γ and IL-4 responses in BMDMs. **a** Time course of transcription in BMDMs after stimulation with 10 ng/ml LPS+10 ng/ml IFN-γ or 100 ng/ml IL-4 for selected targets measured by RT-qPCR (mean ± SEM, n=3). **b** Heatmaps of response in BMDMs after stimulation for 24 h with 10 ng/ml IFN-γ + a range of LPS doses combined with a range of doses of IL-4. Data represent *Arg1* and *Nos2* transcript levels measured by RT-qPCR and Chi3l3, IL-6, IL-12p40, and TNFα protein levels measured by ELISA (mean, n=3). *P*-value represents the significance of the interaction between the stimulations calculated by a 2-way ANOVA. **c** Phospho-Stat1 and phos-pho-Stat6 expression by flow cytometry in BMDMs after stimulation with 10 ng/ml LPS+10 ng/ml IFN-γ, 100 ng/ml IL-4, or the combination for 30 min. **d** Density scatter plots of scRNA-seq transcript counts across individual cells for *Rsad2* vs. *Ifit2* (top) and *Dok2* vs. *Cflar* (bottom). ρ indicates Spearman correlation coefficient. Gene expression is shown as ln(transcript count +1).

**Supplementary Fig. 2 (associated with Fig. 2).**
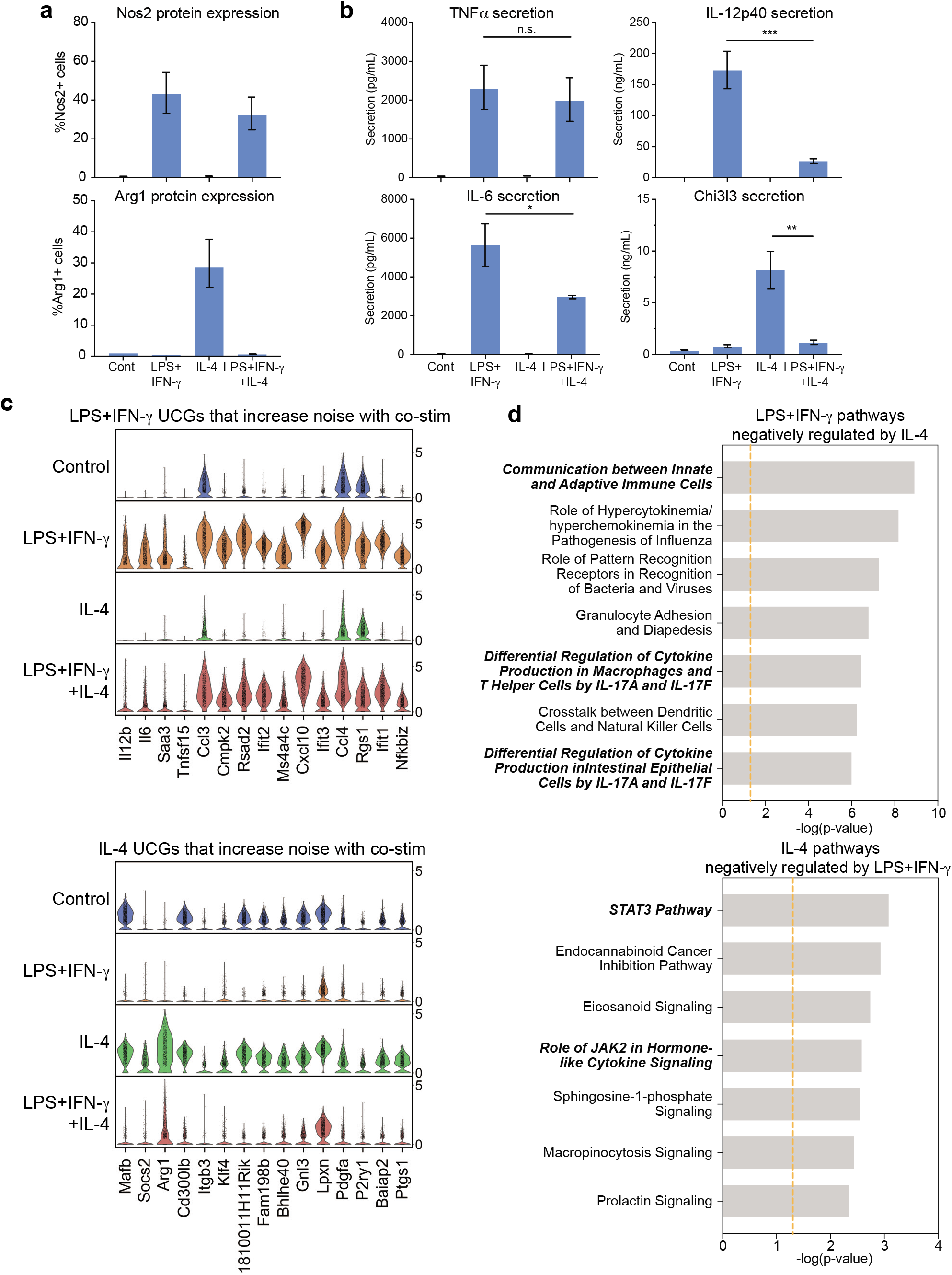
Cross-inhibition and cell-to-cell noise in gene expression after co-stimulation. **a** Levels of Nos2 (top) and Arg1 (bottom) protein detected by flow cytometry after stimulation for 6 h with 10 ng/ml LPS+10 ng/ml IFN-γ, 100 ng/ml IL-4, or the combination (mean ± SEM, n=2). **b** Levels of TNFα, IL-12p40, IL-6 and Chi3l3 secretion after stimulation for 24 h with 10 ng/ml LPS+10 ng/ml IFN-γ, 100 ng/ml IL-4, or the combination (mean ± SEM, n=3). Sidak test, *\*p* ≤ 0.05, \*\**p* ≤ 0.01, \*\*\**p* ≤ 0.001. **c** Violin plots of scRNA-seq measurements of LPS+IFN-γ-induced (top) and IL-4-induced (bottom) genes with the highest change in cell-to-cell gene expression noise measured by Fano factor. **d** Result from Ingenuity Pathway Analysis listing the top 7 canonical pathways that are associated with the set of LPS+IFN-γ-induced (top) or IL-4-induced (bottom) genes that are inhibited after co-stimulation. *P*-values indicate the significance of the overlap between the canonical pathway and the inhibited genes.

**Supplementary Fig. 3 (associated with Fig. 4).**
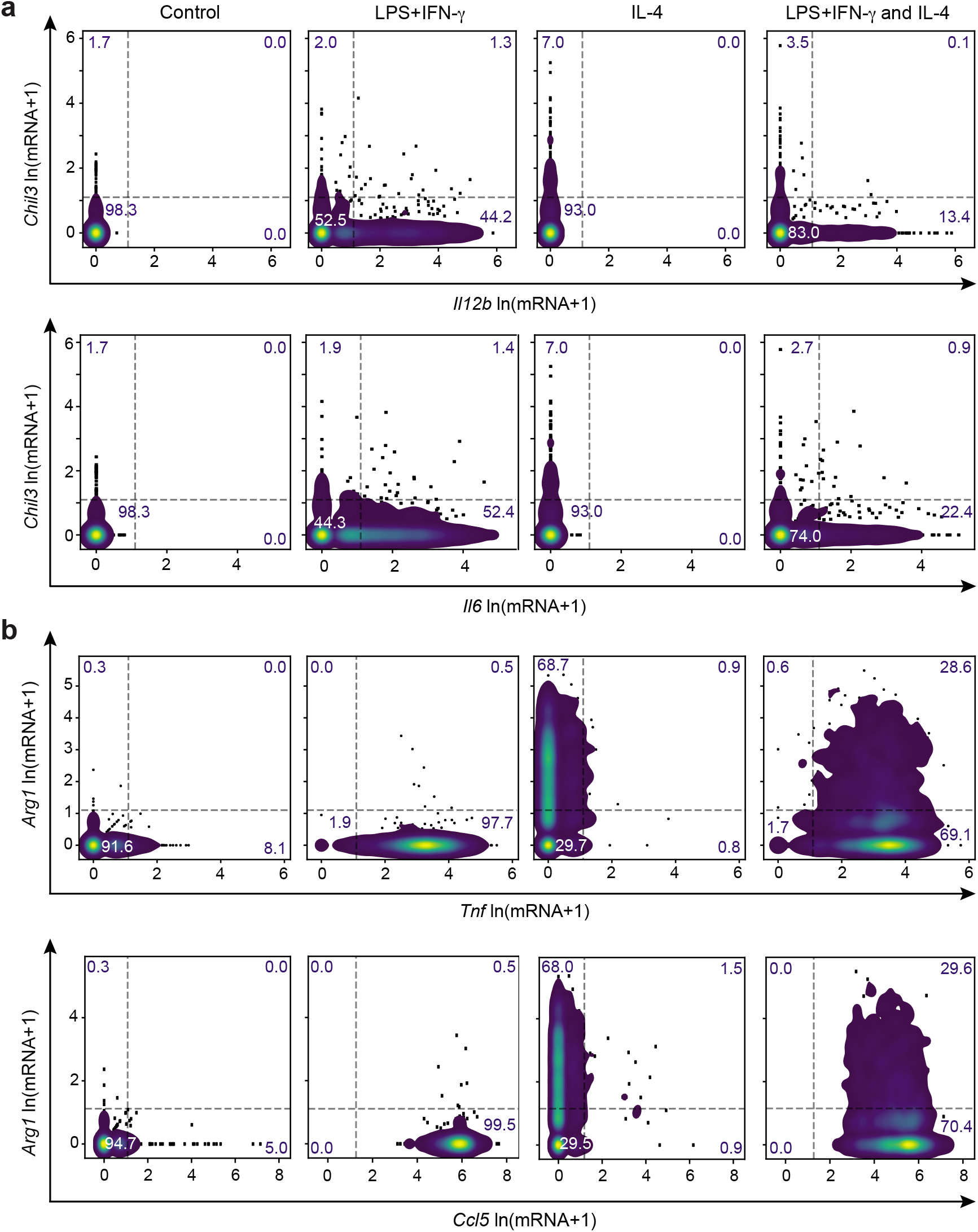
Orthogonal expression is observed for *Il6* and *Il12b* with *Chil3* but not for *Tnf* and *Ccl5* with *Arg1.* Density scatter plots of scRNA-seq transcript counts across individual cells for *Chil3* vs. *Il12b* (**a**, top) and *Il6* (**a**, bottom); and for *Arg1* vs. *Tnf* (**b**, top) and *Ccl5* (**b**, bottom). Data is represented as ln(transcript count +1).

**Supplementary Fig. 4 (associated with Fig. 5).**
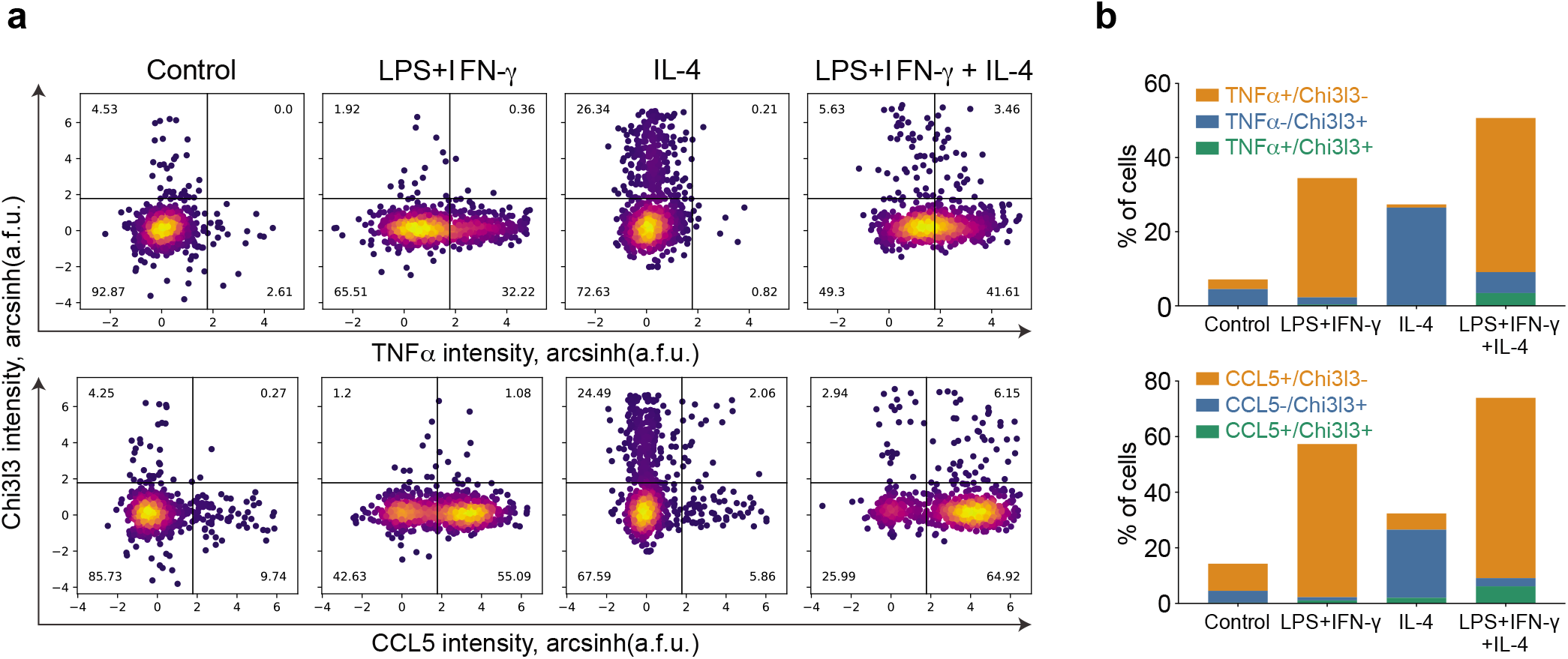
TNFα and CCL5 are not orthogonally secreted with Chi3l3. A multiplexed single-cell secretion assay was used to measure cytokine/chemokine production in individual BMDMs stimulated for 48 h with media alone, 10 ng/ml LPS+10ng/ml IFN-γ, 100 ng/ml IL-4, or both. **a** Density scatter plots of single-cell secretion intensity (a.f.u.) across individual cells for Chi3l3 vs TNFα (top) and Chi3l3 vs CCL5 (bottom). **b** Quantification of single-positive and double-positive cells after 48-hour co-stimulation for secretion of Chi3l3 and TNFα (top) and Chi3l3 and CCL5 (bottom). Data is pooled from 2 biological replicates.

